# Clade 2.3.4.4b H5N1 high pathogenicity avian influenza virus (HPAIV) from the 2021/22 epizootic is highly duck adapted and poorly adapted to chickens

**DOI:** 10.1101/2023.02.07.527270

**Authors:** Joe James, Elizabeth Billington, Caroline J Warren, Dilhani De Sliva, Cecilia Di Genova, Maisie Airey, Stephanie M. Meyer, Thomas Lewis, Jacob Peers-Dent, Saumya S. Thomas, Abigail Lofts, Natalia Furman, Marek J. Slomka, Ian H. Brown, Ashley C. Banyard

## Abstract

The 2021/2022 epizootic of high pathogenicity avian influenza (HPAIV) remains one of the largest ever in the UK, being caused by a clade 2.3.4.4b H5N1 HPAIV. This epizootic affected more than 145 poultry premises, most likely through independent incursion from infected wild birds, supported by more than 1700 individual detections of H5N1 from wild bird mortalities. Here an H5N1 HPAIV, representative of this epizootic (H5N1-21), was used to investigate its virulence, pathogenesis and transmission in layer chickens and pekin ducks, two species of epidemiological importance. We inoculated both avian species with decreasing H5N1-21 doses. The virus was highly infectious in ducks, with high infection levels and accompanying shedding of viral RNA, even in ducks inoculated with the lowest dose, reflecting the strong waterfowl adaptation of the clade 2.3.4.4 HPAIVs. Duck-to-duck transmission was very efficient, coupled with high environmental contamination. H5N1-21 was frequently detected in water sources, serving as likely sources of infection for ducks, but inhalable dust and aerosols represented low transmission risks. In contrast, chickens inoculated with the highest dose exhibited lower rates of infection compared to ducks. There was no evidence for experimental H5N1-21 transmission to any naive chickens, in two stocking density scenarios, coupled with minimal and infrequent contamination being detected in the chicken environment. Systemic viral dissemination to multiple organs reflected the pathogenesis and high mortalities in both species. In summary, the H5N1-21 virus is highly infectious and transmissible in anseriformes, yet comparatively poorly adapted to galliformes, supporting strong host preferences for wild waterfowl. Key environmental matrices were also identified as being important in the epidemiological spread of this virus during the continuing epizootic.

## INTRODUCTION

Following the emergence of A/goose/Guangdong/1/1996 (GsGd) lineage H5N1 highly pathogenic avian influenza virus (HPAIV) in 1996 in China, this lineage spread to other Asian countries, evolving into diverse clades classified according to the H5 Haemagglutinin (HA) gene phylogeny [1]. Since 2009, several Neuraminidase (NA) gene reassortants have occurred in East Asia among GsGd clade 2.3.4.4 viruses, including the H5N8 subtype first introduced into Europe during autumnal wild waterfowl (anseriforme) migration in 2014 (clade 2.3.4.4a) [2]. Since 2014, there have been repeat incursions of HPAIV into the United Kingdom (UK) poultry sector, driven by the incursion of infected wild bird species on migratory pathways, and typically following a seasonal pattern, peaking during the winter period [3]. The European 2014 H5N8 HPAIV (H5N8-14) was responsible for a mild epizootic in the UK and Europe, whereas the re-emergence of a clade 2.3.4.4b European 2016 H5N8 (H5N8-16) caused a larger, more widespread epizootic over the winter of 2016-17 [4]. Two other relatively localised European epizootics have occurred, one in December 2017 caused by a novel European H5N6 HPAIV (H5N6-17) reassortant [5], and the other in 2019, caused by a virus related to the 2016 European H5N8 HPAIV which re-emerged (H5N8-19) in Central and Eastern Europe [6].

In the 2020/21 autumn/winter season, the AI situation escalated considerably; the UK and European continent as whole experienced the largest HPAIV epizootic at the time, caused primarily by H5N8 HPAIV (H5N8-20) [7, 8]. Within this epizootic, four other H5 HPAIV subtypes (H5N1, H5N3, H5N4, H5N5), and at least 19 distinct genotypes were observed [8, 9]. The H5N1 subtype detected during this period contained the HA and M gene segment genetically related to the H5N8-20, but the remaining gene segments were likely donated from indigenous AIVs circulating in wild bird populations in Europe [8, 10]. A genetically highly related H5N1 was detected in a series of mass-die-off events in great skuas (*Stercorarius skua*) off the coast of Scotland during the summer of 2021 [11]. The multi-subtype epidemiological scenario then changed during 2021/22 with the dominance of H5N1 HPAIV (H5N1-21), with only a small number of other subtypes (H5N2 and H5N8 [a single detection of H5N8 in a wild bird in 2021]) detected [8, 12]. In the UK, the 2021/22 epizootic began with H5N1 isolated from a chicken at a backyard poultry premise (IP) notified on the 24^th^ October 2021 [13]. This virus was an H5N1 HPAIV which shared high genetic similarity to the H5N1 circulating in wild birds at low levels in the previous 2020/21 epizootic [8, 10] and exhibited genetic drift which was evident during its persistence in UK wild birds during summer 2021 [11]. Since this first detection in poultry, there were over 145 confirmed IPs with HPAIV H5N1 in poultry and captive birds across Great Britain (GB) until the defined start of the 2022/23 epizootic on 1^st^ October 2022 [8, 13]. As well as numerous poultry incursions, there were over 1,700 dead wild birds testing positive for H5N1 across GB during this period [13]. These unprecedented infection levels were widely mirrored across continental Europe [7, 8].

The high infection pressure in the two recent epizootics (2020/21 and 2021/22) has resulted in several detections of HPAIV in non-avian wildlife species, likely through opportunistic scavenging behaviour on dead, infected wild birds [3, 14-17]. In additional to sporadic infection of mammalian wildlife species, there has been a human infection reported in the UK [18, 19]. Alongside this unprecedented emergence and spread in the UK and Europe, the H5N1-21 HPAIVs have also spread to North America and onwards to South America following the migration routes of wild birds [20, 21]. The altered seasonal epidemiology of H5N1 HPAIV was again reflected by persistence in UK wild birds over the summer of 2022, with continued poultry incursions also being reported [22].

Together, these observations signify the importance of developing a better understanding of these H5Nx HPAIVs from an avian perspective. The increased incidence of cases during the 2021/22 epizootics reflects a distinct change in the dynamics of infection across Europe. Studies have previously demonstrated *in vivo* that the earlier H5Nx clade 2.3.4.4 HPAIVs are highly anseriforme adapted [23-25]. It is speculated that the H5N1-21 may have acquired yet further increased fitness in anseriformes compared to previous European H5Nx epizootic viruses, so providing an enhanced waterfowl reservoir which then drives poultry outbreaks.

To assess the *in vivo* fitness of the H5N1-21 HPAIV, two species of high importance, ducks as anseriformes, representing a surrogate for wild aquatic birds and domesticated waterfowl poultry, and chickens as galliformes, representative of the poultry species of major commercial importance were assessed. The infectivity, viral tropism, pathogenicity and transmissibility in both these species using an HPAIV H5N1-21, representative of the clade 2.3.4.4b H5N1 from the early part of the 2021/2022 epizootic was investigated. The infectivity and pathogenesis were assessed by inoculating groups of ducks and chickens with three decreasing doses of H5N1-21; intraspecies transmission efficiency in chickens and ducks was also explored in different experimental scenarios. Potential transmission routes were also interrogated by examining environmental contamination with H5N1-21 originating from infected birds.

## METHODS

### Virus origin and propagation

A/chicken/England/053052/2021 (H5N1) (GISAID accession number EPI_ISL_9012457) was isolated from brain tissue of chickens confirmed to have HPAIV H5N1 infection using specified pathogen free (SPF) 9-day-old embryonated fowls eggs (EFE) as described previously [26]. This viral isolate was representative of the 2021/21 HPAIV UK epizootic (Fig. S1). A/chicken/England/053052/2021 (H5N1) represents the first HPAIV detection during the UK 2021-22 epizootic which was notified on 24^th^ October 2021 and is hereby referred to as H5N1-21. H5N1-21 was further propagated in EFEs and titrated in FEFEs to determine the 50% egg infectious dose (EID_50_) [24]. The full genome was sequenced using whole genome sequencing (WGS), as previously described [24], to confirm that no amino-acid polymorphisms had emerged following passage, when compared to the original clinical sequence. Virus stocks were diluted in 0.1M pH 7.2 phosphate buffered saline solution (PBS) for all *in vivo* infections.

### Ethics and Safety Statement

The *in vivo* study was reviewed and approved by the local Animal and Plant Health Agency (APHA) Animal Welfare and Ethical Review Body to comply with the relevant UK legislation, in accord with the UK Home Office (HO) Project License PP7633638. According to the UK’s Advisory Committee on Dangerous Pathogens (ACDP) and the Specified Animal Pathogens Order (SAPO); the GsGd lineage H5Nx HPAIVs are classified at ACDP Hazard Group Level three pathogens and SAPO level four, hence all animal and laboratory work involving infectious material was conducted using licensed containment level (CL) 3 facilities at APHA [23-25]. Welfare monitoring of up to three times daily assessed the birds for humane endpoints following the onset of clinical disease, enabling any decisions to be made concerning the need for euthanasia. All birds had access to food and water *ad libitum*.

### Birds

High health status layer chickens (Hy-Line Brown), and Pekin ducks (Cherry Valley hybrid) of mixed sex (52% female, 48% male; and 51% female, 49% male respectively) were sourced from Hy-Line UK Ltd. (UK) and Cherry Valley Farms Ltd. (UK), respectively. Birds were obtained at one-day age and reared until 3 weeks-of-age at time of infection. Both species were acclimatised for seven days prior to the first procedures which included swabbing (oropharyngeal (Op) and cloacal (C)) and bleeding from a superficial vein prior to inoculation. These samples were tested using an M-gene RRT-PCR [27] and serological testing [28], the latter using the ELISA kit (ID Screen® Influenza A Antibody Competition Multi-species, IDVet, France).

### Experimental design: 50% minimum infectious dose (MID_50_) for ducks and chickens

To investigate the 50% minimum infectious dose (MID_50_) and viral pathogenicity in both species, three groups of chickens (n=6/group) and ducks (n=6/group) were inoculated with 100 μl of H5N1-21, administered via the oculo-nasal route, containing either a low (10^3^ EID_50_), medium (10^4^ EID_50_), or high (10^5^ EID_50_) dose of the H5N1-21 virus (Fig. S1). Chickens were housed in Perspex pens 120 (l) x 180 cm (w) (area of 2.16 m^2^). Ducks were housed in Perspex pens 120 (l) x 240 cm (w) (area of 2.88 m^2^). Both duck and chicken pens contained bell gravity fed water drinkers and feed hoppers, while each duck pen contained a 1 m^2^ pond. From 1 to 14 days post infection (dpi), Op and C swabs were collected daily from all birds. Three feathers of an immature character, and between 3-5 cm maximum length (including the calamus which associated with the adjacent skin follicle; [29, 30]), were collected from the abdominal area of individual birds on alternate days from 1 dpi until 14 dpi. To minimize the risk of birds experiencing severe clinical signs due to HPAIV infection, birds were euthanised upon reaching a defined humane endpoint, while any surviving birds were culled at study-end (14 dpi), when terminal bleeds collected via heart puncture. Opportunistic *post-mortems* were performed on a sub-set of birds which were euthanised or found dead during infection. Environmental samples were collected daily until 10 dpi from each high dose pen. The dose required to cause MID_50_ was calculated as previously defined [24].

### Experimental design: intraspecies transmission among chickens and ducks

Transmission in chickens was assessed by housing the chickens at low or high stocking densities. For the low stocking density experiment, transmission was assessed for each of the doses (low [10^3^ EID_50_], medium [10^4^ EID_50_], and high [10^5^ EID_50_] per chicken) by simply extending the MID_50_ study. The six directly infected chickens in each group were referred to as the donor (D0) chickens (Fig. S1B). Six naive “first recipient” (R1) chickens were introduced for co-housing into each of the three dose groups at 6 hrs post-infection. The 18 co-housed chickens were therefore held at a low bird density of 0.18 m^2^ / chicken (Fig.S1B). Sampling was extended to include the R1 chickens, as described for the MID_50_ study.

For the high stocking density experiment, an increased number of 15 D0 chickens were inoculated with a high dose of H5N1-21 (10^5^ EID_50_ per chicken) via the oculo-nasal route, as described for the MID_50_ study (Fig. S2). At six hours post infection, 15 R1 chickens were introduced for co-housing with the D0 chickens, with all 30 housed in a Perspex pen (120 (l) x 180 cm (w)), at a density of 0.072 m^2^ / chicken.

Transmission in ducks was assessed by modelling a realistic scenario of an initial AIV incursion into a farmed anseriforme premises or in a wild bird setting, where a small number of D0 ducks were co-housed with a larger number of R1 ducks. Three D0 ducks were directly inoculated with a medium dose 10^4^ EID_50_ per duck of the H5N1-21, via the oculo-nasal route (Fig. S2). All 21 ducks were housed in a large, open pen design 340 cm (w) x 310 cm (l) (10.54 m^2^ floor area), with access to a 2.55cm^2^ pond (density of 0.58 m^2^ / duck).

### Clinical scoring and monitoring

All birds were observed and clinically scored three times a day (morning, afternoon, and evening). A range of clinical signs were monitored, including visual, behavioural, and neurological signs which have weighted scores based on their severity (1-7) (Table S2). To prevent clinical deterioration resulting in severe clinical signs, birds were euthanised upon reaching the humane endpoint (≥7 cumulative clinical score/timepoint) as described previously [23-25]. For ethical reasons, any birds housed singularly, due to euthanasia or death of others within the group, were also euthanised.

### Clinical sample collection and processing

Swabs were taken from the Op and C cavity. Swabs were individually cut and placed into 1 ml of Leibovitz’s L-15 Medium (Gibco; (LM) [25]), and the supernatant used for RNA extraction or stored at -80 °C until further use. During *post-mortem* examination, approximately 50 mg of tissue from selected organs was similarly collected into 1 ml LM and mechanically homogenised, with the homogenate used for RNA extraction. For virus-specific immunohistochemical (IHC) analysis, thin tissue sections from organ samples were fixed in 10% (v/v) buffered formalin and embedded in paraffin for investigation by standard Haemoxylin and Eosin (H and E) staining. In addition, influenza A virus type-specific IHC staining, using a broadly reactive anti-NP monoclonal antibody, was undertaken, with the overall intensity of virus-specific staining in a given tissue assessed by a semi-quantitative scoring system (from 0 [no staining] to 4 [high intensity staining]) [30]. Blood was collected prior to infection though superficial vein bleeds, or at study end through cardiac puncture under terminal anaesthesia. The serum was separated from whole blood and used in serological analyses.

### Environmental sample collection and processing

Environmental samples were collected from all *in vivo* experiments. In the duck and chicken pens, when one air sampler was used, it was placed in a central position within the pen. In contrast where multiple air samplers were used, they were placed on top of the bell drinker, food hopper, and in a central position within the pen (over the pond for ducks). The actual air sampler receptacle was placed out of the reach of the birds and did not come into direct contact with any bird for the duration of the study. The samplers consisted of Gelatine Filters (25 mm, with a 3 μm pore size) housed in a Button Aerosol Sampler (SKC Ltd.), with each connected to an APEX2 air pump (Casella). Air samples were collected at indicated time points prior to, and following infection, whereby air was drawn for 6 hours at a flow rate of 2.00 litres/min (total volume of 720 litres). While the filter has a nominal pore size of 3.0 μm, these filters have a higher capture efficiency of sub-micron particles through inertial impaction and diffusional interception [31], thereby enabling immobilisation of inhalable particles (size range: <1 μm – 100 μm) which include a range of bioaerosols and inhalable dust. After each 6-hour collection, the gelatine filters were dissolved in 2 ml LM, and total RNA was extracted from the supernatant.

Pond water (ducks only), drinking water, straw and faecal samples were collected from the pens of all birds at indicated timepoints, as described previously [24]. For the straw and faecal material, samples from two different areas of the pen (‘front’ and ‘back’) were pooled, and approximately 1 g of material was added to 2 ml of PBS. RNA was extracted from the supernatants obtained from the solid matrices and directly from the water samples.

### RNA extraction and AIV reverse transcription Real-Time PCR (RRT-PCR)

RNA was extracted from LM obtained from swabs, tissues, feathers and environmental samples by using the MagMAX™ CORE Nucleic Acid Purification Kit (Thermo Fisher Scientific™) as part of the robotic Mechanical Lysis Module (KingFisher Flex system; Life Technologies), according to the ‘manufacturer’s instructions. 2 μl volumes of extracted RNA were tested by the M-gene RRT-PCR using the primers and probes designed by Nagy, Černíková [27]. RRT-PCR Ct values < 36 were considered AIV positive, sub-threshold values in the range Ct 36.01-39.99 and Ct 40 (“No Ct”) were interpreted as negative [32]. A ten-fold dilution series of titrated H5N1-21 HPAIV RNA was used to construct a standard curve using Agilent Aria software (Agilent, UK) to determine PCR efficiency, with Ct values obtained from viral specimens converted to relative equivalent units (REUs/ml) by correlation with the EID_50_/ml values of the extracted viral standard curve as previously described [33]. Birds were considered to have been infected with H5N1-21 based on (i) positive RRT-PCR results, indicating shedding of viral RNA (vRNA)), from at least two swabs (Op or C) or (ii) one M-gene RRT-PCR positive swab in addition to seroconversion by either Hemagglutination inhibition assays (HAI) or FluA ELISA. Absence of viral RNA (vRNA) shedding, as evidenced by negative swab results throughout the monitoring period, and serological negativity (if available), classified a bird as uninfected.

### Serological analysis

Hemagglutination inhibition assays (HAI) were conducted to determine the presence of antibodies for specific H5N1 subtypes according to standard methods [28]. Whole blood samples were centrifuged for 5 minutes at 2000 rpm to separate the serum from the blood cells. Sera were incubated at 56 °C for 30 minutes to inactivate complement. The duck sera were pre-absorbed by adding packed chicken red blood cells and incubated in at 4 °C overnight to prevent non-specific agglutination. The beta-propiolactone inactivated viral antigen [34] A/Chicken/Wales/053969/21 (H5N1) (GISAID Accession number EPI_ISL_9012618), representative of the antigenic profile of the 2021/22 H5N1 epizootic, was used at four HA units. Reciprocal HAI titres of greater than 1/16 were considered seropositive [23]. All sera were also tested by the multi-species influenza A ELISA (IDEXX, France) and the FluA ELISA (IDVET, France) according to the manufacturer’s instructions.

### Statistical analysis

Area under the curve (AUC) analysis was performed on the shedding profile of individual birds using GraphPad Prism v8. The values for each individual bird were calculated as REUs/ml. A mean was calculated from the individual bird AUC values, with birds exhibiting no shedding obtaining a value of 0 REUs/ml. Data was compared for significance in GraphPad Prism v8 using one-way ANOVA and multiple comparisons tests. Statistical significance was reported on p-values ≤0.05.

## RESULTS

### MID_50_ determination and accompanying mortality: Clade 2.3.4.4b 2021 H5N1 is more infectious for ducks than chickens

To investigate the virulence of the HPAIV 2021/2022 H5N1 epizootic in galliformes and anseriformes, chickens and ducks were infected with three different doses of H5N1-21 (low [10^3^ EID_50_], medium [10^4^ EID_50_], or high [10^5^ EID_50_]). Among the ducks, all three doses resulted in all (100%) of the ducks becoming infected, as evidenced the H5N1-21 shedding (Fig. 1A-F). Successful H5N1-21 infection of all six ducks in the low dose group (Fig. 1A-F) demonstrated the high viral infectivity in this host, with the MID_50_ being <10^3^ EID_50_. By contrast, in the chickens which received the same three doses of H5N1-21, infection was apparent in none (0%), one (17%) and four of six (67%) in the low, medium, and high dose groups, respectively (Fig. 1G-L), corresponding to a greater MID_50_ of 10^4.67^ EID_50_ in chickens. H5N1-21 infection in both species resulted in mortality, which included decisions to euthanize all 18 ducks as they reached humane clinical endpoints (Fig. 2A, Table S1). In each of the three dose groups, five of the six ducks were euthanized on welfare grounds because they had registered individual clinical scores in the range 7-10 (Table S1, Table S2), with the intervention applied to prevent these 15 ducks from experiencing clinical disease *in extremis*. The neurological signs which contributed to the decision to euthanize included frequent observation of loss of balance and tremors among these 18 infected ducks, although torticollis and / or other more moderate or mild signs (lethargy and conjunctivitis were most common) (Table S1, Table S2). The mean death times (MDTs) for the ducks euthanized on clinical grounds was 7.2, 5.6 and 4.3 days in the low, medium, and high dose groups respectively. Welfare considerations also stipulated that the single remaining duck in each group could not remain housed alone, so these three infected ducks were also euthanized in each group, despite displaying lower clinical scores (Table S1, Table S2).

**Fig. 1.**
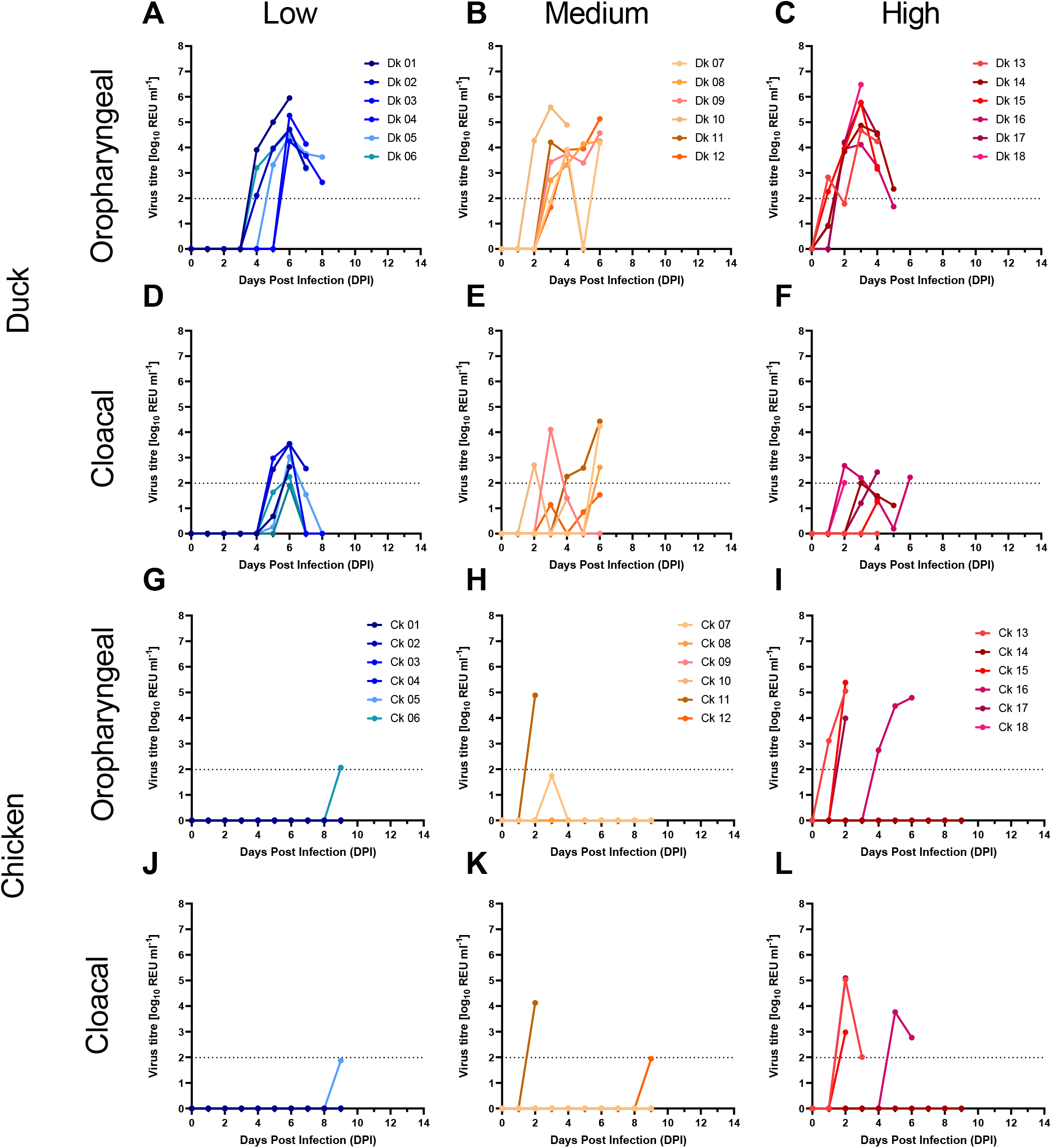
Virus shedding from ducks and chickens which were directly infected with different doses of H5N1-21 HPAIV. Shedding was measured for the Op (A-C and G-I) and cloacal (D-F and J-L) samples in ducks (A-F) and chickens (G-L), following infection with H5N1-21 at low (10^3^ EID_50_) (A, D, G, J), medium (10^4^ EID_50_) (B, E, H, K) and high (10^5^ EID_50_) (C, F, I, L) doses. Viral titres were determined by using the M-gene RRT-PCR. Dotted horizontal lines indicate the M-gene RRT-PCR positive cut-off.

**Fig. 2.**
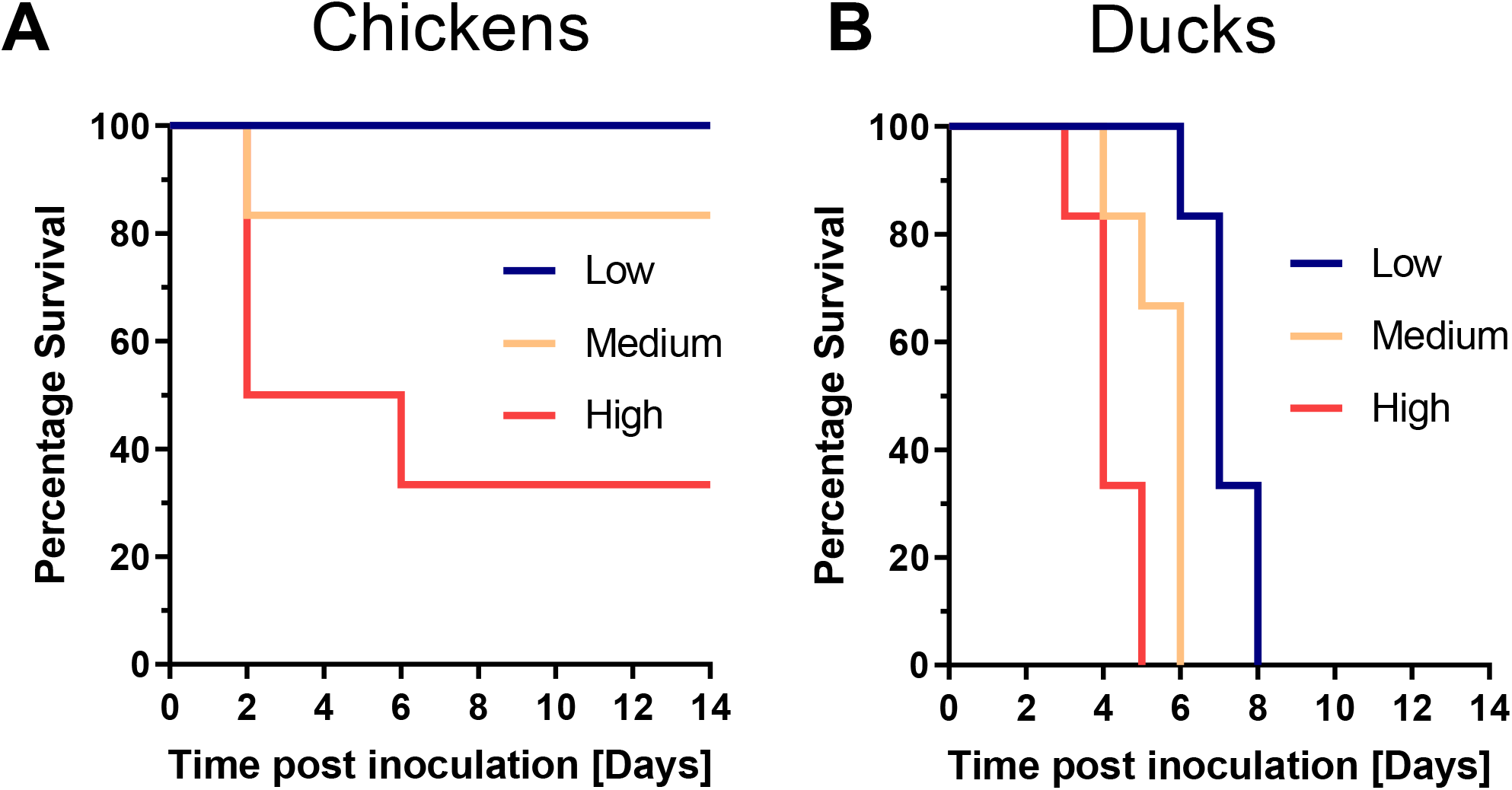
Survival of chickens and ducks following infection with H5N1-21 HPAIV. Percentage survival of (A) chickens and (B) ducks infected with H5N1-21. Birds were each infected via the oculo-nasal route with 100 μl inoculum containing either a low (103 EID50), medium (104 EID50) or high (105 EID50) doses of H5N1-21. Birds were monitored three times daily and scored for clinical signs. Mortalities among chickens and ducks included both euthanised and “found dead” birds, with all ducks in (B) having been euthanised due to clinical signs (Table S1). Consequently,(B) does not include mortality of the final surviving duck in each dose group because these were culled to comply with welfare guidelines concerning single housing of birds.

Although H5N1-21 was less infectious in chickens compared to ducks, infection in chickens always resulted in death (Fig. 2A), with an MDT of 3.1 days in the high-dose chicken group which was shorter than in the corresponding high dose duck group (Table S1). A single infected chicken in the medium-dose group also succumbed to infection rapidly at 2 dpi. Three of the five chicken deaths were characterized by a rapid clinical deterioration, as no obvious clinical signs had been observed at earlier timepoints (Table S1). However, the other two chicken mortalities were welfare-based euthanasia interventions guided by increasing clinical signs which most frequently included loss of balance, lethargy, and closed eyes (Table S2).

Additional examination of H5N1-21 vRNA shedding revealed that all 18 of the directly infected ducks shed H5N1-21 vRNA from the Op cavity in all dose groups, with peak shedding titres between 10^4^ and 10^7^ REUs/ml (Fig. 1 A-C). Onset of H5N1-21 vRNA shedding was delayed as the dose decreased, with vRNA being shed from the Op cavity at 1 dpi, 2 dpi and 4 dpi for the low, medium, and high dose, respectively (Fig. 1 A-C). Cloacal shedding of vRNA was considerably less pronounced than Op vRNA shedding in ducks, with equivalent titres of less than 10^5^ REUs/ml being sporadically detected (Fig. 1 D-F). although these differences between total Op and C shedding (AUC values) was only statistically significant for the high dose inoculated groups (p-value <0.0001). In the five H5N1-21 infected chickens, the shorter period of vRNA shedding included Op shedding in all five birds, but C shedding was only detected in four (Fig. 1 G-L). Absence of H5N1-21 infection in all surviving chickens was confirmed by seronegative results, as measured by HAI or influenza A anti-NP ELISA (Fig. S3A).

### H5N1-21 pathogenesis in ducks and chickens

H5N1-21 tissue tropism was investigated among the infected birds which had died or been euthanised during the MID_50_ experiment, with total RNA extracts tested by M-gene RRT-PCR from a range of tissues obtained from three chickens and six ducks (Fig. 3, Table S1). vRNA was detected in a wide range of tissues from the medium and high dose chickens, except for the pancreas, spleen, kidney and brain (Fig. 3A). All three chickens displayed the highest vRNA titres in the lung, caecal tonsil, proventriculus and feather calamus, with mean titres of 3.88×10^5^, 2.87×10^5^, 2.75×10^5^, and 5.51×10^4^ REUs/ml, respectively. Two of three chickens also displayed very high vRNA titres in the heart, with a mean of 2.11×10^6^ REUs/ml (Fig. 3A). In the ducks which had been infected at all three doses, high H5N1-21 vRNA titres were again detected in a wide range of tissues except for the spleen and kidney. High vRNA titres were detected in the feather calami and the brain of all but one duck, with mean vRNA titres of 2.34×10^6^, and 2.82×10^7^ REUs/ml, respectively (Fig. 3A).

**Fig. 3.**
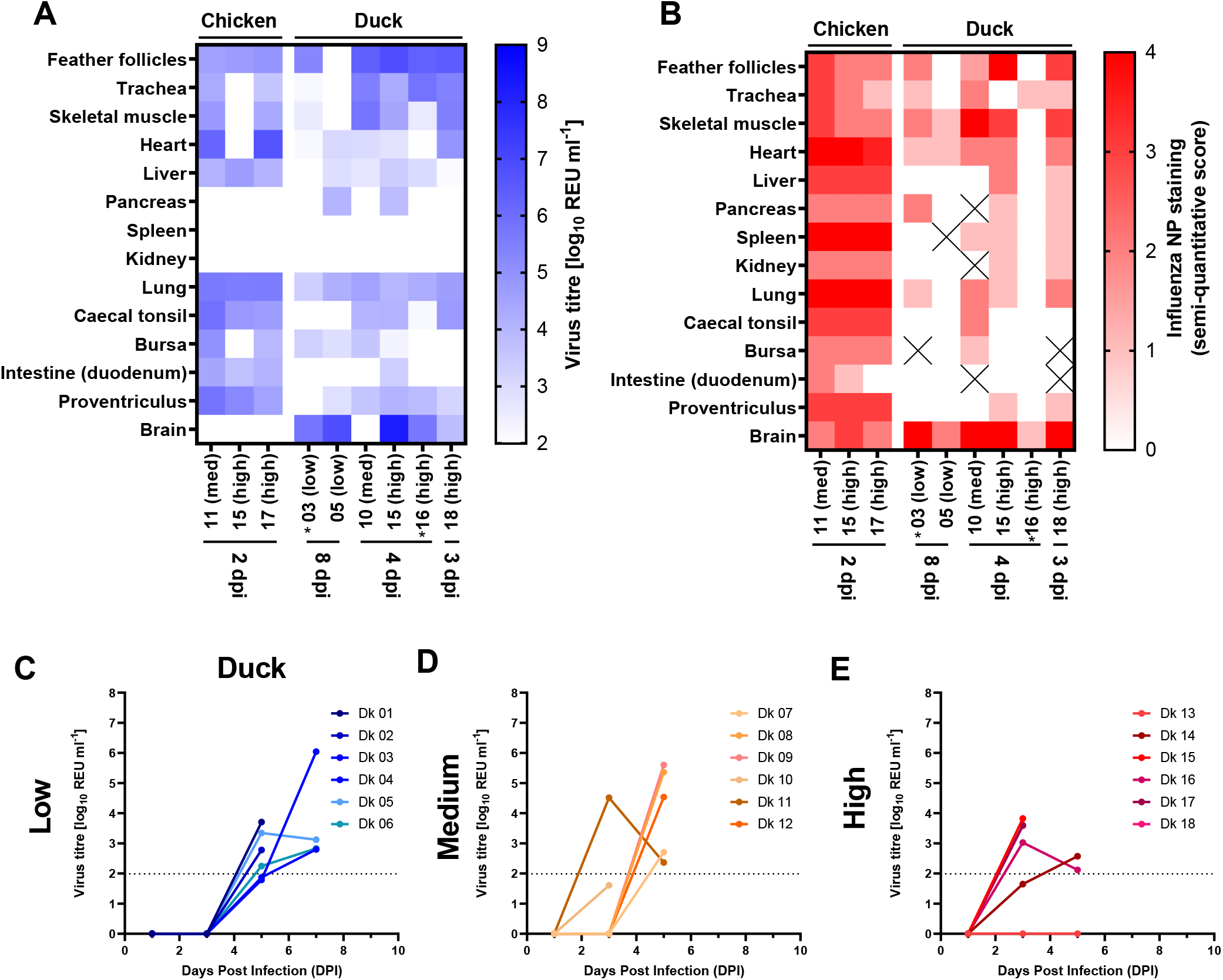
Organ tropism in chickens and ducks directly infected with different doses of the H5N1-21 HPAIV. M-gene RRT-PCR determined vRNA titres (A) or influenza NP semi-quantitative staining from IHC (B) from tissues taken from chicken and duck mortalities, infected low (10^3^ EID_50_), medium (med) (10^4^ EID_50_) and high (10^5^ EID_50_) doses of H5N1-21 HPAIV. Asterix indicates birds euthanised due to single housing, and not because of severe clinical signs. A ‘cross’ in the heat map indicates samples could not be tested. **C-D**. vRNA titres were similarly determined in the feather follicles from ducks infected with the same three doses of H5N1-21. Horizontal dotted lines indicate the positive cut-off.

IHC analysis was conducted on sections obtained from the same birds and tissues used for vRNA analysis (Fig. 3B). The presence of the influenza A nucleoprotein (NP) was detected in almost all tissue sections collected from the three chickens, except for an absence of NP staining in the intestine of one chicken (Fig. 3B). NP staining was evident in the pancreas, spleen, kidney and brain tissues of the three infected chickens, despite vRNA not being detected (Fig. 3A, B). The chicken heart, spleen and lung tissues had the highest levels of NP staining, whereas the pancreas, kidney, bursa, and intestine exhibited the lowest abundance of NP staining (Fig. 3B). In the ducks, NP staining was more sporadically detected across the tissue panel, and generally at lower abundance compared to respective tissues from the chickens. NP staining was consistently detected in the brain of all six ducks, at relatively high abundance, consistent with high vRNA titres also found in the brain of these ducks (Fig. 3A, B). NP staining was also evident in the skeletal muscle and heart tissue in five out of the six ducks, and in the feather follicle, trachea, and lung in four out of the six ducks (Fig. 3B).

vRNA titres were determined in the feather calami as a measure of H5N1-21 dissemination. vRNA and NP staining was detected in the feathers collected from the three chicken mortalities at 2 dpi (Fig. 3A, B), all of which had been shown to be H5N1-21 infected by virtue of vRNA shedding (Fig. 1G-L). Feathers were also collected from the remaining chickens at all dose groups (low, medium and high) at 3, 5 and 7 dpi, although none of these feathers were positive for vRNA (data not shown). Feathers were similarly collected from infected ducks in the three dose groups between 1-7 dpi, which revealed the presence of H5N1-21 vRNA (Fig. 3C-E). A dose-effect was apparent in that the vRNA detected in the feather, which was first detected at 3 dpi in three ducks in the high-dose group, but in only one duck at this time point in the medium-dose group, while the earliest detection in the low-dose group was in five ducks at 5 dpi (Fig. 3C-E).

### H5N1-21 is highly efficient at transmitting between ducks but not chickens

We investigated transmission of H5N1-21 between chickens, at two different stocking densities (low and high) (Fig. S1, Fig. S2). The Low stocking density transmission experiment was carried out as an extension of the above chicken MID_50_ study, where the three directly inoculated chicken groups served as the D0 chickens, with six R1 chickens introduced for co-housing 6 hours after D0 infection. In the low stocking density transmission experiment, there was no evidence of any H5N1-21 transmission between chickens following inoculation with the three doses (Fig. S4). None of the contact (R1) chickens displayed any clinical signs or seroconverted by HAI or anti-NP ELISA (Fig. S3A).

To investigate transmission a at higher stocking density, a greater number of co-housed chickens (15 D0 and 15 R1 chickens) were used (Fig. S2A). To ensure over 50% of the directly inoculated chickens were infected, the high dose inoculum was used (10^5^ EID_50_ per chicken). H5N1-21 vRNA was detected in swab samples in 93% (14/15) of the D0 chickens starting from 2 dpi, where each chicken registered at least two instances of positive shedding, or positive shedding from both tracts (Fig. 4A, B). Fourteen out of 15 D0 chickens which became infected displayed severe clinical signs and were either found dead or euthanised by 2 dpi (11 found dead; 3 euthanised) with an MDT of 1.98 days (Fig. 4A, B, Table S2). Despite successful infection of the D0 chickens, none of the 15 in -contact R1 chickens became infected (Fig. 4A, B), which was confirmed by the absence of clinical disease and lack of seroconversion by HAI or anti-NP ELISA (Table S2, Fig. S3B).

**Fig. 4.**
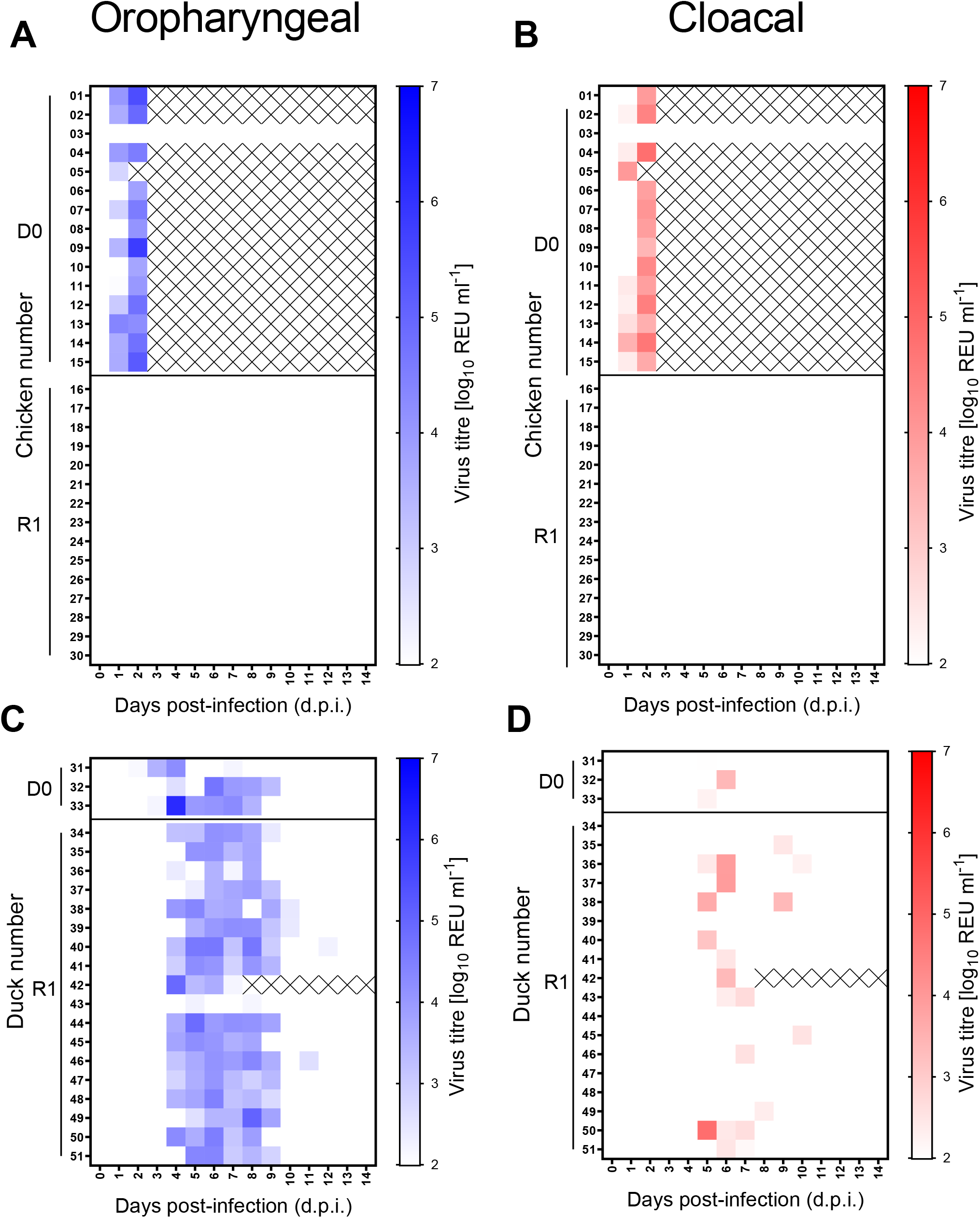
H5N1-21 shedding during transmission attempts from directly-infected to contact birds (chickens and ducks). Virus RNA titres from directly infected (D0) or contact (R1) chickens (A and B) and ducks (C and D) infected via intra nasal and intra ocular route with 10^5^ EID_50_ (chickens) or 10^4^ EID_50_ (ducks) of H5N1-21. Contacts (R1) were introduced 6 hours after infection. vRNA titres were determined by using the M-gene RRT-PCR derived from swabs taken from the Op (A and C; blue) or cloacal cavities (B and D; red) and displayed graphically as a heat map. White indicates no Ct or REU values below the limit of positivity. Shaded colours represent higher vRNA titres which were positive by the M-gene RRT-PCR. A ‘cross’ indicates that no sample was collected at the specific time point due to the bird being euthanised following severe clinical signs.

Transmission dynamics was also investigated in ducks as a surrogate for spread among wild anseriformes, or for wild waterfowl mediated incursions into domesticated anseriforme poultry. Experimental modelling included a more natural outdoor scenario for wild bird mediated incursion of infection, where only three D0 ducks were directly infected in a larger area than used in the chicken transmission scenario. In addition, the larger area provided low density duck housing (density of 0.58 m^2^ / duck) with a greater number of introduced R1 ducks (Fig. S2B). All three D0 ducks became infected, with vRNA detected at positive levels in Op swabs in 100% (3/3) of the D0 ducks, with initial onset of Op vRNA shedding evident in the range 2-4 dpi for these three ducks (Fig. 4C). Cloacal shedding was sporadic, with vRNA only being detected on two days in 66% (2/3) of the D0 ducks (Fig. 4D). However, no statistically significant differences were observed between total Op and C vRNA shedding (AUC values), or between total vRNA shedding (AUC values) from D0 compared to R1 ducks (data not shown). Transmission was 100% efficient, with all 18 R1 ducks shedding vRNA from Op or C swabs on multiple days (Fig. 4C, D). All surviving ducks seroconverted by HAI and anti-NP ELISA (Fig. S3B). By examining the initial onset of vRNA shedding among the R1 ducks, 67% (12/18) became infected by 4 dpi (2 days following the onset of shedding in the D0 ducks), with the remaining 33% (6/18) becoming infected by 5 dpi (Fig. 4C, D).

### Environmental contamination from infected and contact ducks is greater than from chickens

Environmental specimens were collected daily during the MID_50_ and low-stocking density transmission determination in chickens which were infected with the high dose of H5N1-21 (Fig. S1B). Only one bedding sample collected at 7 dpi contained vRNA in the low stocking scenario at a relatively low vRNA titre (2.95×10^3^ REUs/ml), while all the other environmental specimens were negative (Fig. 5A). Detection of H5N1-21 vRNA in the bedding at 7 dpi correlated with high Op and C shedding from one chicken (Fig. 1I and L; 5A), although earlier contamination of the bedding could not be excluded. Similarly, environmental contamination in the high chicken stocking scenario (Fig. S2A) was also minimal, with vRNA being detected in only a single drinking water sample at 5 dpi, again at a low vRNA titre (4.01×10^3^ REUs/ml) (Fig. 5B).

**Fig. 5.**
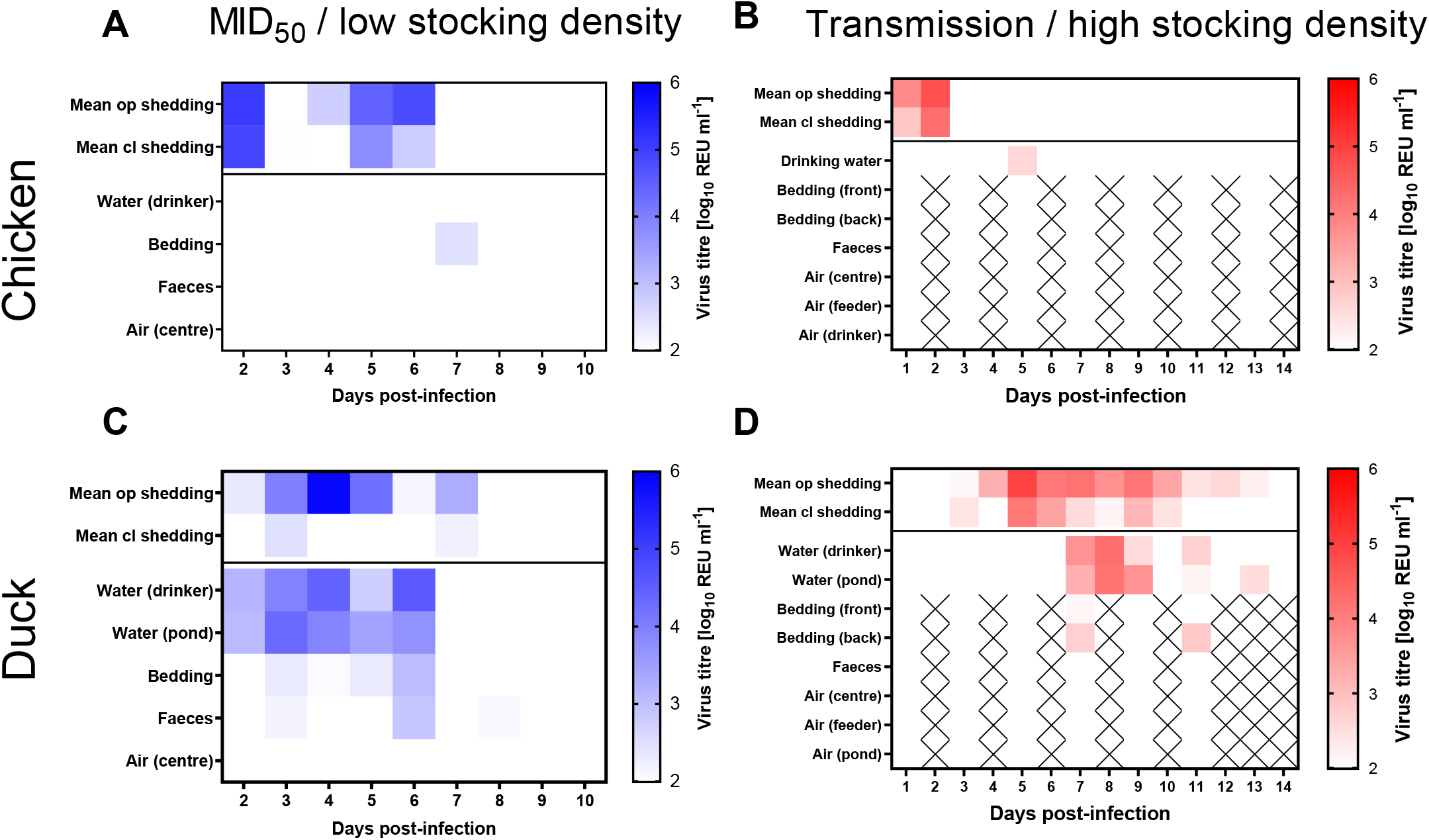
Environmental contamination with H5N1-21 HPAIV. Virus RNA titres were determined in environmental samples collected from the areas which housed chickens at (A) low and (B) high stocking densities. Environmental specimens were similarly collected and tested from the areas which housed ducks at (C) the MID_50_ and (D) transmission study. Viral titres were determined by the M-gene RRT-PCR and are displayed graphically as a heat map, with white indicating “No Ct” or REU values below the positive cut-off. Shaded colours represent positive vRNA titres. ‘Cross’ indicates that no sample was collected at the specific time point. The top two rows on all graphs represent the mean oropharyngeal (op) and cloacal (cl) shedding titres from all birds, obtained by combining the individual bird shedding data shown in Fig. 1 and 4 from the two experimental designs.

By contrast, H5N1-21 vRNA contamination of the environment was detected much more frequently in the duck experiments conducted at both stocking densities (Fig. S1A, S2B). vRNA was detected continually in both the drinking and pond water between 2-6 dpi in the MID_50_ study (Fig. 5C). Maximal titres of 3.96×10^4^ REUs/ml and 2.51×10^4^ REUs/ml were attained in the drinking and pond waters, respectively; detection in water correlated with vRNA shedding (predominantly OP) which was observed in these ducks (Fig. 1C, F, 5C). vRNA was also detected in the bedding and faecal samples at lower titres at a single timepoint (6 dpi), with titres of 1.10×10^3^ REUs/ml and 7.83×10^2^ REUs/ml, respectively (Fig. 5C). The infected ducks in the transmission study also contaminated both sources of water, with H5N1-21 Vrna being detected between 7-13 dpi (Fig. 5D). This period of water contamination again occurred after the onset of H5N1-21 vRNA shedding in the ducks (Fig. 4C, D, 5D), with the 17 infected R1 ducks being the likely major contributors to the viral contamination of both water sources. Maximal titres of 1.84×10^4^ REUs/ml and 1.51×10^4^ REUs/ml were attained in the drinking and pond waters, respectively (Fig. 5D). Low-level contamination of bedding was observed on only two sampling days (7 and 11 dpi) in this duck housing, with vRNA titres of 5.24×10^2^ REUs/ml and 6.78×10^2^ REUs/ml, respectively (Fig. 5D).

Air samples were collected daily in both the chicken and duck experiments at both stocking densities by using samplers and filters capable of collecting inhalable bioaerosols and airborne fomites (e.g., inhalable dust). No vRNA was detected above the positive threshold level in any of the air samples collected from the four housing settings at any time-point post infection (Fig. 5A-D).

## DISCUSSION

Since autumn 2014, there have been seasonal incursions of clade 2.3.4.4 H5Nx HPAIVs into poultry sectors across Europe mediated by wild birds [3, 5-8, 35, 36]. An extensive HPAIV clade 2.3.4.4 epizootic has affected the UK and Europe during 2020-21 being caused primarily by the H5N8 subtype, with an even more extensive and impactful epizootic now being caused by H5N1 since autumn 2021 [7, 8]. Key underlying questions concern the identification of virological factors which are responsible for the extensive and sustained nature of the currently continuing H5N1 epizootic. As such, a series of *in vivo* investigations were designed to assess the viral infectivity, transmissibility and pathogenesis in chicken and duck hosts.

Determination of the MID_50_ in ducks underlined the highly infectious nature of H5N1-21 in this host. Ducks served as an experimental surrogate to demonstrate how readily H5N1-21 may be sustained in both migratory wild waterfowl and farmed anseriformes. In East Asia, both domesticated and wild anseriformes have been central to the maintenance of the GsGd HPAIVs for over 20 years [37], with the clade 2.3.4.4 H5Nx HPAIVs having been subsequently disseminated to other continents where epizootics of varying magnitudes and duration have occurred [20, 36]. The MID_50_ for H5N1-21 was determined as being <10^3^ EID_50_, and similar in ducks as earlier UK H5N8 (2016) and H5N6 (2017) clade 2.3.4.4 HPAIVs [24, 25], as well as for a Danish-origin H5N8 (2016) [38].

The H5N1-21 demonstrated a strong waterfowl adaptation by its efficient transmission from three directly infected (R1) ducks to 18 contact (D0) ducks, so affirming the prior intra-species transmission findings in ducks noted for earlier clade 2.3.4.4 UK isolates, namely H5N8 (2014) and H5N6 (2017), plus the Danish H5N8 (2016) isolate [23, 24, 38]. A striking observation included the very rapid transmission of H5N1-21 from three D0 ducks inoculated with a 4 log_10_ EID_50_ dose (initial Op vRNA shedding observed between 2 and 4 dpi in the D0 ducks), resulting in 12 infected R1 contact ducks by 4 dpi, and the remaining six R1 ducks acquiring infection by 5 dpi. Infection kinetics were also assessed in D0 ducks inoculated with the same dose of the earlier clade 2.3.4.4 H5N8 (2014), with the first onset of vRNA shedding in D0 ducks observed at 2 dpi and all seven live D0 ducks acquiring infection by 5 dpi [23]. Rapid H5N8 (2014) transmission followed to the R1 ducks, with six of eight and all eight R1 ducks infected at 3- and 4-days post-introduction for cohousing, respectively. Although the numbers of D0 and R1 ducks in these transmission studies differed, both H5N1-21 and H5N8 (2014) demonstrated fast and efficient spread among this species [23]. By comparison, a 10^4^ EID_50_ dose of the earlier UK H5N6 (2017) appeared to initiate direct infection (via shedding of infectious material) in three D0 ducks slightly more slowly (first Op vRNA shedding observed at 3 dpi), followed by all five R1 contacts acquiring infection by 13 days after introduction for cohousing [24], indicating a much slower transmission of infection compared to H5N1-21 and H5N8 (2014). These findings suggest although different clade 2.3.4.4 H5Nx HPAIVs spread efficiently from infected to naïve ducks, the transmission kinetics of individual strains does appear to differ, and that this is a virus-specific characteristic which may be reflected in how rapidly a particular clade 2.3.4.4 strain may disseminate within wild waterfowl, as opposed to the extent of its spread among waterfowl. Although epidemiological and other environmental factors may also contribute to clade 2.3.4.4 kinetics and related spread within wild waterfowl, the rapid transmission from three D0 to 18 R1 ducks observed for H5N1-21 may be mirrored in the largest ever number of clade 2.3.4.4 wild bird cases in the UK which were dominated by anseriformes during winter 2021-22 [8, 10] with an increased detections in Charadriiformes (shorebirds) over the summer of 2022 [11]. Consequently, a greater degree of ongoing infection may impact more quickly within wild waterfowl populations prior to resolution of shedding and seroconversion, which may increase the frequency of transmission events at the wild bird and terrestrial poultry interface [23-25].

Compared to ducks, MID_50_ determination in chickens revealed a higher value (10^4.67^ EID_50_), indicating a lower viral infectivity when H5N1-21 was inoculated directly into these two species. Because H5N1-2020 is a very similar direct ancestor of H5N1 from 2020 (H5N1-20) [8], the dose range applied in the current H5N1-21 MID_50_ experiment was informed by the outcomes of the earlier H5N1-2020 MID_50_ experiment in layer chickens of the same age which resulted in 100% infection and mortality following inoculation with 6 log_10_ EID_50_ of H5N1-2020 [39]. Another MID_50_ study also underlined that clade 2.3.4.4 H5Nx HPAIVs are more infectious in ducks compared to chickens [38], with similar observations having also been made for earlier GsGd H5N1 HPAIVs (clade 2.2) [40]. Such *in vivo* comparisons of quantified MID_50_ values in different avian species have served as an indicator of a given AIV strains adaptation to terrestrial poultry [41, 42]. In addition, in the current study, two stocking densities were used to assess chicken-to-chicken transmission of H5N1-21, by using equal numbers of D0 and R1 chickens. The areas available per chicken were selected to reflect the maximum stocking densities which are mandated for commercial poultry in the UK broiler and layer sectors [43-45]. Despite the successful infection of all D0 chickens in both experiments, there was no evidence of any transmission to the R1 contacts. The higher stocking density transmission attempt also included a greater number of chickens (15 D0 and 15 R1 chickens). Leyson, Youk [38] similarly observed failed transmission of the Danish clade 2.3.4.4 H5N8 (2016) among chickens, although smaller total numbers of chickens were cohoused in their study.

Clearly, the lack of chicken-to-chicken transmission for H5N1 clade 2.3.4.4 contradicts the UK poultry epidemiology where a significant proportion of commercial chicken premises have been affected by the ongoing H5N1 HPAIV epizootic, with a high percentage of infected chickens recorded during outbreak investigations at clinically affected epidemiological units [46]. These field observations suggest that factors such as, the larger numbers of chickens with greater environmental contamination at commercial premises and prolonged contact with infected carcasses, along with other possible local or environmental factors, serve to successfully initiate the spread of infection within this species. One instance of successful H5N8 (Hungarian isolate, 2017) clade 2.3.4.4 transmission featured 20 D0 and 20 R1 chickens [47]. However, no humane endpoint was used, because the authors believed that HPAIV transmission would be affected by the earlier removal of strongly shedding and diseased chickens. This approach differed from the considerations for the current study’s experimental design, where any infected birds which were beginning to display severe clinical signs were promptly euthanised.

The contrast between the efficient and absent transmission of H5Nx clade 2.3.4.4 HPAIVs in ducks and chickens, respectively, has been previously noted in studies which compared successful intra-species transmission among ducks with inter-species transmission from ducks to chickens, which included introduction of subsequent chicken groups. D0 ducks were successfully directly infected with H5N8 (2014), and indeed the first chicken contacts (R1) were all successfully infected [23]. However, introduction of subsequent naïve chickens resulted in onward H5N8 (2014) transmission in this species becoming increasingly less efficient [23]. Seekings, Warren [24] demonstrated that D0 ducks infected with H5N6 (UK 2017 isolate) were responsible for frequently detected environmental contamination with this virus, particularly in the pond and drinking water, and that this contamination coincided with shedding from both D0 and contact infected R1 ducks. By contrast, transmission among chickens of H5N8 and H5N1 clade 2.3.4.4 HPAIVs (both isolated in late 2020) was inefficient and was accompanied by an absence of environmental contamination which included bedding, faeces and drinking water [39]. This finding suggested that the occasional transmission to R1 chickens was likely due to close contact between the chickens.

In the current study, efficient duck-to-duck transmission of H5N1-21 was accompanied by clear viral contamination in the co-housing environment, again most notably in the water samples where H5N1-21 detection coincided with viral shedding from the ducks. For the failed low and high stocking density H5N1-21 transmission in chickens, we used the same layer chicken breed which featured in the studies of Seekings, Warren [39], but with only single instances of relatively low-level viral contamination in the bedding and drinking water in the low and high stocking density transmission attempts, respectively. In summary, for the numbers of birds which featured in the H5N1-21 *in vivo* experiments, it appears that the high degree of viral contamination of drinking and pond water is a key factor in enabling the efficient duck-to-duck transmission. By contrast, the minimal environmental contamination with H5N1-21 observed in the chicken housing would appear to have been insufficient to result in any viral transmission to cohoused R1 chickens, while close contact of naïve R1 with infected D0 chickens was insufficient to result in any instances of transmission. The failure of any H5N1-21 transmission was observed even when 15 D0 and 15 R1 chickens were cohoused at high density, with initial inoculation having been carried-out with the dose which was gauged from the MID_50_ findings in the low stocking density H5N1-21 experiment in chickens, together with the MID_50_ and associated chicken to chicken transmission findings with H5N1-2020 [39]. It is probably that the rapid MDT for the D0 chickens is not an adequate time for transmission to the R1 contacts, as hypothesised for earlier H5N1 GsGd HPAIVs among chickens [48].

A novel aspect of the study was to include air sampling in all the H5N1-21 *in vivo* experiments to assess potential transmission due to microscopic particles which include biological droplets, aerosols or airborne fomites (such as inhalable dust). These particles may harbour infectious virus, with aerosols typically being less than 5 μm in diameter [49-51]. In a field environment, infectious H5N8 20/21 has been detected in the air sampled on infected commercial poultry premises, with larger particle sizes (such as inhalable dust) harbouring higher titres of virus [52]. However, in the case of the H5N1-21 infected chickens and ducks in the current study, there was no evidence of any airborne viral presence. Environmental contamination, which includes contaminated water and fomites, has been long suspected as being a source of AIV infection at poultry premises and persistence for onward spread [53]. While there are obvious virological differences between the field study and the data presented here, it should be stressed that this study was conducted in a high containment animal facility under constant negative air pressure (25 room air volume changes per hour), as such, a lower concentration of particles is likely to be present in the air [54]. vRNA contamination in water samples underlined its highly likely role in significantly contributing to the highly efficient and rapid transmission among ducks. It must be emphasised that the pond and drinking waters were replaced and refreshed daily, hence the levels of H5N1-21 vRNA detection represented accumulation during the preceding 24 hours. It has also been demonstrated that deliberate experimental contamination of water with clade 2.3.4.4 H5N8-2016 isolates from the UK and The Netherlands can result in successful infection of naïve chickens [55], with similar observations having been made for mallards [56]. By specifically identifying how environmental contamination features in the mechanism of H5Nx clade 2.3.4.4 HPAIV transmission, this study along with others has confirmed the importance of biosecurity to prevent water contamination at poultry premises from becoming a source of infection and onward spread [53, 57].

The H5N1-21 infected chickens and ducks displayed differing mortality patterns. In both the low and high stocking density transmission attempts among chickens, all infected chickens died, this being a typical clinical outcome for any HPAIV infection in this important commercial galliforme species [58, 59]; a mixture of chickens which were found dead and some which required euthanasia due to severe clinical signs was observed. A dose effect was apparent; a shorter MDT was associated with higher infecting dose (Fig 2A), with the 10^5^ EID_50_ dose administered in the high stocking density revealing a similarly rapid MDT which, as noted above, may have been insufficiently long to enable effective shedding of infectious material and subsequent transmission to R1 contact chickens. Systemic dissemination of H5N1-21 antigen and vRNA was apparent among the chicken mortalities, as consistently observed in previous infections in chickens with European H5Nx clade 2.3.4.4 HPAIV [23, 24, 38, 39, 60, 61].

A similar observation was made for the ducks in the MID_50_ experiment which all required euthanasia, albeit at a longer MDT when the same H5N1-21 doses were compared in both species. However, other than the culling of the final surviving ducks in each dose group for welfare reasons, all the earlier decisions to euthanise ducks were prompted by the observation of clear neurological signs. Both H5N1-21 vRNA and antigen were frequently detected in the brains of infected ducks, with neurotropism a likely contributor to the aggressive pathogenesis observed in ducks, albeit dissemination of H5Nx GsGd HPAIVs to other organs has been previously observed to contribute to duck mortality [30, 62] including the earlier H5N8 clade 2.3.4.4 isolates from 2016 [25, 61]. These studies, along with those of others, reveal a wide range of different mortality outcomes for Pekin ducks experimentally infected with different European H5Nx clade 2.3.4.4 HPAIVs, with age being a factor likely predicting reduced clinical disease [23, 24, 38, 55, 63]. Seroconversion and decline in viral shedding in ducks is an indicator of a resolved infection, with clearance of systemic infection from organs reported for H5N6-2017 infected ducks [24]. Investigations H5N1 HPAIV incursions into UK commercial ducks during 2022 have included flocks where ducks were shown to be both shedding vRNA and / or had seroconverted [46]. Investigations H5N1 HPAIV incursions into UK commercial ducks during 2022 have included flocks where ducks were shown to be both shedding vRNA and / or had seroconverted [46].

It was observed that H5N1-21 vRNA was absent in the spleens and kidneys obtained from all sampled mortalities in both species, although viral antigen was in both organs of ducks and chickens.

Discrepancies in detecting viral tropism by the two methods in some tissue specimens may be explained by non-uniform distribution of H5N1-21 infection within a particular organ, as suggested previously [23]. Strong virus-specific IHC staining of the chicken spleens may have reflected the lymphocytic accumulation of viral antigen, as opposed to H5N1-21 replication or accumulation of genomic RNA-containing viral particles. Feather-associated H5N1-21 vRNA was observed in both ducks and chickens, and may be considered an indicator of systemic infection, as noted previously for European H5Nx clade 2.3.4.4 in experimental infections in both species [23-25]. Feathers deposited at in vicinity of outbreak locations also pose a potential risk for subsequent HPAIV infection of naïve poultry [64].

Clear differences were observed in clinical outcomes between the H5N1-21 infected ducks in the duck MID_50_ determination and duck transmission experiments. The former ducks included many decisions to euthanise based on neurological clinical signs, while during the latter experiment all the ducks displayed at least some clinical signs, only one required euthanasia due to its severity. Differences in mortality outcomes in infected Pekin ducks have been observed previously for H5Nx clade 2.3.4.4 viruses when comparing *in vivo* experiments conducted independently at different institutions, as in the case of European H5N6 (late 2017) HPAIVs where 60% [55] and 7% [24] deaths were reported. Factors which may have affected the strikingly different mortality outcomes in these H5N6-infected ducks may have included age, direct inoculation and / or contact exposure routes to attain infection, plus possible strain variation between the two H5N6-2017 isolates from the UK and The Netherlands. In addition, local differences such as the housing environment and welfare considerations to euthanise may have also been influenced by an element of subjective opinion concerning clinical severity. The successful detection of seroconversion in ducks following efficient and rapid transmission of H5N1-21 reaffirmed that serological surveillance may remain important for identifying current or previous instances of incursion into domesticated anseriformes, particularly where overt clinical signs may not be frequently apparent at commercial duck premises [24].

At the time of writing, in early 2023, the clade 2.3.4.4 epizootic initiated by the H5N1-21 HPAIV is continuing to cause poultry outbreaks and wild bird cases in the UK, Europe and the Americas [13], underlying the timeliness of the investigations of infectivity, transmission, environmental contamination and pathogenesis which we have compared in chickens and ducks. The H5N1-21 strain evolved from the H5N1 HPAIV which circulated as a minor population in the 2020/21 epizootic (H5N1-20) after acquiring the HA and M gene segments from the contemporary H5N8-20 HPAIV which was epidemiologically dominant at the time [8, 10]. In addition to its frequent detection in UK poultry and wild birds, this strain has also caused a human infection [8, 18]. However, as the epizootic has progressed, a variant with five amino acid differences in the HA has emerged and largely replaced this strain [10]. In addition, several other variants have emerged, and have been detected in both poultry outbreaks and wild birds sporadically [8]. These variants contain several internal gene segments closely related to those detected in indigenous European wild bird AIVs [8, 10]. The impact of these novel HPAIVs in terms of infectivity, pathogenicity and transmissibility in different avian species remains to be explored, particularly as extensive dissemination within the wild bird reservoir appears to have been a major determinant of the magnitude of this ongoing H5N1 clade 2.3.4.4 epizootic. The fitness H5N1-21 in ducks, as evidenced by the infectivity (MID_50_) and transmission findings, has demonstrated a strong adaptive advantage in anseriformes, leading to continuing circulation and opportunities for further viral evolution, including reassortment events which may serve to make the clade 2.3.4.4 epizootic more complex and diverse.

The repeated incursions of H5Nx clade 2.3.4.4 viruses globally demonstrate a continuing threat to the poultry industry with its economic consequences. Concerns have also arisen concerning the ecological and biodiversity impact of H5N1 HPAIV upon particular seabird populations, as observed along the UK coast [22]. Strong viral adaptation to wild waterfowl which includes migratory species has featured in the inter-continental spread of the current H5N1 HPAIV [21], underlying the global nature of this problem. It is difficult to predict the next variant to emerge from among the clade 2.3.4.4 H5Nx HPAIVs. Therefore, it is important to characterize the phenotype of newly emerging strains promptly, to help inform potential mitigations and interventions which will control or limit the current or any subsequent clade 2.3.4.4 epizootic.

## Funding information

This work was supported by the Biotechnology and Biological Sciences Research Council (BBSRC) and Department for Environment, Food and Rural Affairs (Defra, UK) research initiative ‘FluMAP’ [grant number BB/X006204/1]. Funding was also provided by the Defra and the Devolved Administrations of Scotland and Wales, through SE2213 ‘FLUFUTURES 2’.

## Acknowledgements

The authors would like to thank Daniel Maskell, Nadia Chew, Amy Miller and Elena Mather for collection and processing of samples from the *in vivo* experiments.

## Author contribution

J.J., I.H.B., M.J.S. and A.C.B. conceived and designed the study; J.J., E.B., C.J.W., D.D.S., C.D.G., M.A., S.M., T.L., J.P. S.S.T, A.L., N.F. and M.J.S. conducted experimental work; J.J., E.B., C.J.W., D.D.S., A.L., N.F., and M.J.S. interpreted the data; J.J., M.J.S., E.B., D.D.S., I.H.B and A.C.B. wrote the manuscript.

## Conflict of interest

The authors declare no conflicts of interest.

## Notes

### Competing Interest Statement

The authors have declared no competing interest.

