## Supplementary material for "Clade 2.3.4.4b H5N1 high pathogenicity avian influenza virus (HPAIV) from the 2021/22 epizootic is highly duck adapted and poorly adapted to chickens": Table S1

**Table S1.** $MID_{50}$ experiment: Clinical end points in ducks and chickens following inoculation with different doses of H5N1-21.

| Species | Bird ID | Time of death (dpi) | Clinical score at termination | Euthanised (E) or found dead (FD) | Dose group | Mean death time (MDT) per group^ |
|---|---|---|---|---|---|---|
| Duck | 1 | 6 | 8 | E | Low | 7.2 days |
| | 2 | 7.5 | 9 | E | | |
| | 3 | 8 | 4 | $E^{pm}$* | | |
| | 4 | 7 | 8 | E | | |
| | 5 | 8 | 8 | $E^{pm}$ | | |
| | 6 | 7.5 | 10 | E | | |
| Duck | 7 | 6 | 10 | $E^{pm}$ | Medium | 5.6 days |
| | 8 | 6 | 8 | E | | |
| | 9 | 6.5 | 5 | E* | | |
| | 10 | 4 | 8 | E | | |
| | 11 | 5.5 | 7 | E | | |
| | 12 | 6.5 | 9 | E | | |
| Duck | 13 | 4.75 | 8 | E | High | 4.25 days |
| | 14 | 5 | 9 | E | | |
| | 15 | 4 | 9 | $E^{pm}$ | | |
| | 16 | 5 | 5 | $E^{pm}$* | | |
| | 17 | 4 | 9 | E | | |
| | 18 | 3.5 | 8 | $E^{pm}$ | | |
| Chicken | 11 | 2 | n/a | $FD^{pm}$ | Medium | 2 days |
| | 13 | 2.5 | n/a | FD | High | 3.12 days |
| | 15 | 2 | n/a | $FD^{pm}$ | | |
| | 16 | 6 | 7 | E | | |
| | 17 | 2 | 10 | $E^{pm}$ | | |

* Euthanised due to ethical restrictions which prevent the continued housing of single surviving birds; pm, post mortem analysis conducted; dpi, days post infection; n/a, not applicable; ^ MDT calculations do not include the surviving birds.
