## Supplementary material for "Clade 2.3.4.4b H5N1 high pathogenicity avian influenza virus (HPAIV) from the 2021/22 epizootic is highly duck adapted and poorly adapted to chickens": Table S2

**Table S2.** Frequency of observed clinical signs in ducks and chickens directly inoculated with different doses of H5N1-21 or ducks placed in direct contact.

| | Total counts of clinical sign occurrence | | | | | | | | |
|---|---|---|---|---|---|---|---|---|---|
| | Ducks | | | | | Chickens | | | |
| | $MID_{50}$ experiment | | | Transmission | | MID experiment | | | Transmission |
| **Clinical Sign** | **D0 (low)** | **D0 (medium)** | **D0 (high)** | **D0 (med)** | **R1** | **D0 (low)** | **D0 (medium)** | **D0 (high)** | **D0 (high)** |
| **Huddling/ruffled feathers** | 18 | 17 | 11 | 0 | 3 | 0 | 1 | 8 | 3 |
| **Eyes closed** | 3 | 6 | 3 | 0 | 3 | 0 | 0 | 5 | 3 |
| **Conjunctivitis** | 31 | 21 | 34 | 0 | 13 | 0 | 0 | 0 | 0 |
| **Dropped wings** | 2 | 0 | 0 | 0 | 0 | 0 | 0 | 1 | 2 |
| **Oedema** | 0 | 4 | 1 | 0 | 0 | 0 | 0 | 0 | 0 |
| **Cyanosis** | 0 | 0 | 0 | 0 | 9 | 0 | 0 | 0 | 0 |
| **Lethargy** | 30 | 20 | 21 | 4 | 10 | 0 | 1 | 3 | 2 |
| **Discharge eyes/beak** | 0 | 0 | 2 | 0 | 0 | 0 | 0 | 1 | 0 |
| **Diarrhoea** | 5 | 0 | 2 | 6 | 11 | 0 | 0 | 1 | 1 |
| **Visual reduction in weight** | 0 | 0 | 1 | 0 | 0 | 0 | 0 | 1 | 0 |
| **Loss of balance** | 9 | 10 | 9 | 0 | 0 | 0 | 0 | 4 | 2 |
| **Tremors** | 5 | 4 | 6 | 1 | 1 | 0 | 0 | 1 | 1 |
| **Torticollis** | 0 | 2 | 0 | 0 | 1 | 0 | 0 | 0 | 0 |
| **Paralysis** | 0 | 0 | 0 | 0 | 1 | 0 | 0 | 0 | 2 |
| **Found Dead** | 0 | 0 | 0 | 0 | 0 | 0 | 1 | 0 | 11 |

Clinical signs were not observed in chickens placed in contact (R1) and are therefore no data is shown; $MID_{50}$, 50% minimum infectious dose; D0, directly infected; R1, contact; low, $10^3$ $EID_{50}$; medium, $10^4$ $EID_{50}$; high, $10^5$ $EID_{50}$.
