## Supplementary material for "Clade 2.3.4.4b H5N1 high pathogenicity avian influenza virus (HPAIV) from the 2021/22 epizootic is highly duck adapted and poorly adapted to chickens": Fig. S1

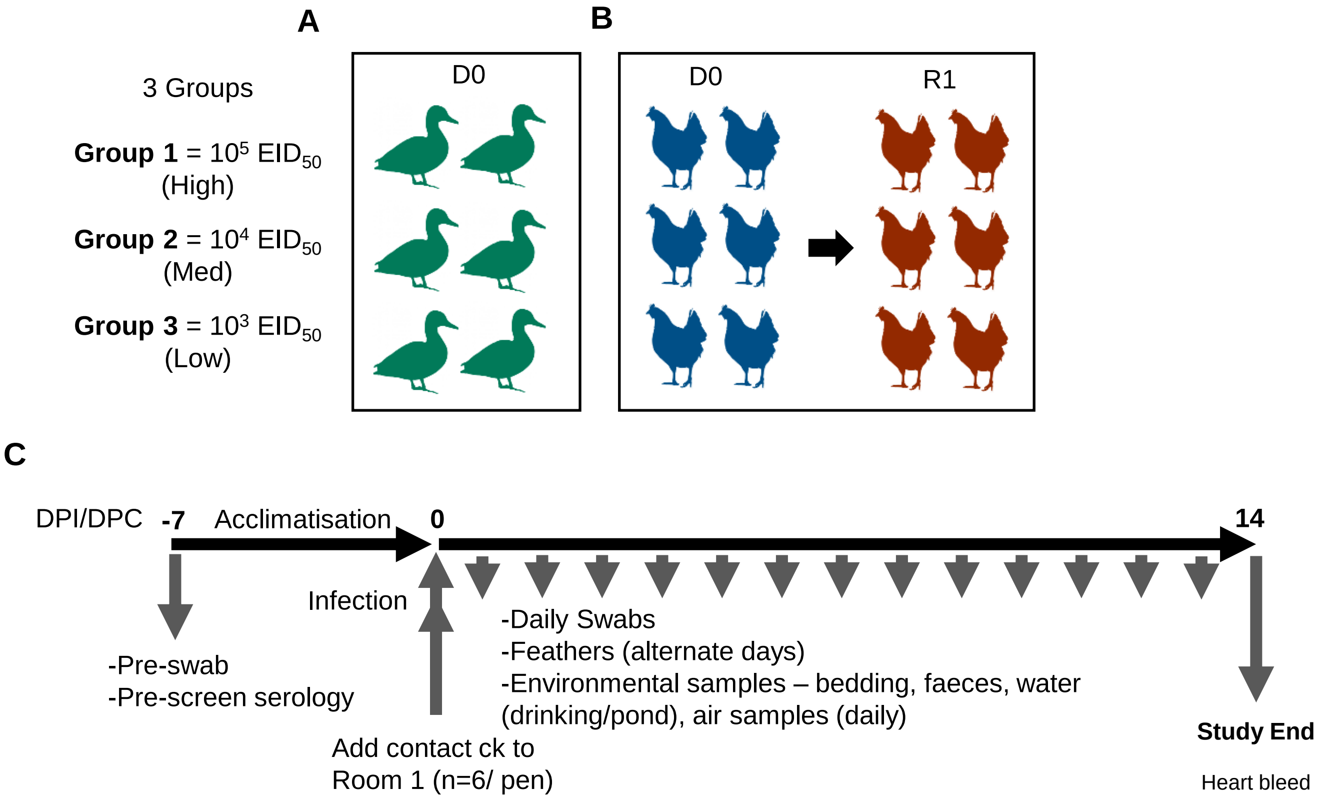


**Fig. S1. Experimental design: 50% minimum infectious dose ($MID_{50}$) and transmission of H5N1-21 HPAIV.**

All birds were housed at a low stocking density. The schematic diagram illustrates the experimental design for (A) ducks and (B) chickens to assess 50% minimum infectious dose ($MID_{50}$) of H5N1 21, with the latter including extension to investigate attempted transmission from D0 to R1 chickens. (C) illustrates the sampling timeline for the experiments.
