## Supplementary material for "Clade 2.3.4.4b H5N1 high pathogenicity avian influenza virus (HPAIV) from the 2021/22 epizootic is highly duck adapted and poorly adapted to chickens": Fig. S2

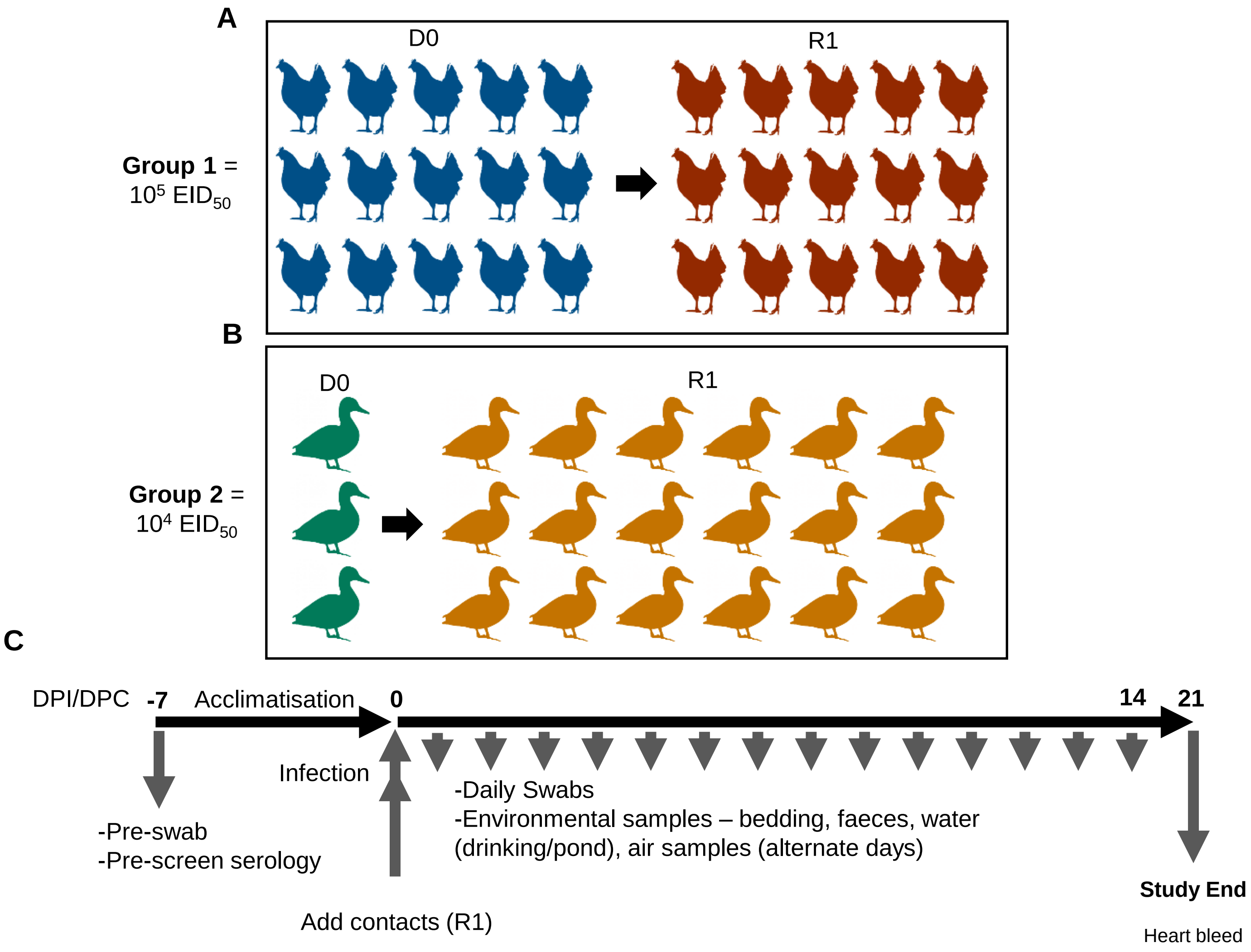


**Fig. S2. Transmission of H5N1-21 HPAIV experimental design.**

Schematic diagram of the experimental design used to assess H5N1 21 transmission dynamics among (A) ducks and (B) chickens housed at a high stocking density. (C) illustrates the sampling timeline for the experiments.
