## Supplementary material for "Clade 2.3.4.4b H5N1 high pathogenicity avian influenza virus (HPAIV) from the 2021/22 epizootic is highly duck adapted and poorly adapted to chickens": Fig. S3

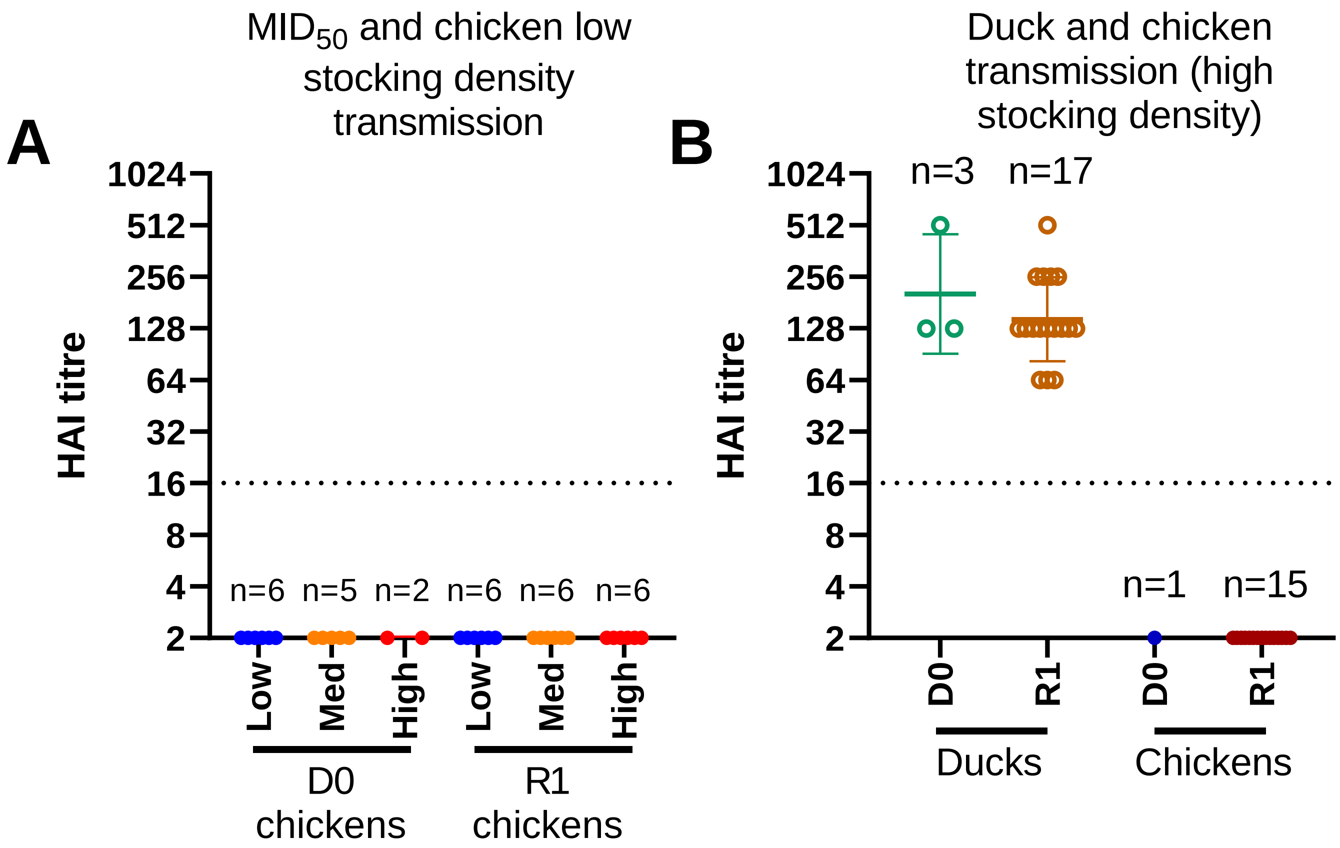


**Fig. S3. Seroconversion of directly infected and contact ducks and chickens following infection and contact exposure to H5N1-21.**

Sera obtained from D0 and R1 ducks and chickens at (A) 14 dpi and (B) 21 dpi. Haemagglutinin inhibition (HI) assays were performed using the selected 2021 H5N1 HPAIV antigen (see Methods). Geometric means of HI titres and their standard deviations are shown. Dotted horizontal line indicates the positive cut-off, with titres ≥1/16 considered positive. Seropositivity was also confirmed using influenza A anti-nucleoprotein (NP) ELISAs; open circles indicate sera positive by both multi-species influenza A ELISA (IDEXX, France) and the FluA ELISA (IDVET, France); solid circles indicate sera negative by both ELISAs.
