## Supplementary material for "Clade 2.3.4.4b H5N1 high pathogenicity avian influenza virus (HPAIV) from the 2021/22 epizootic is highly duck adapted and poorly adapted to chickens": Fig. S4

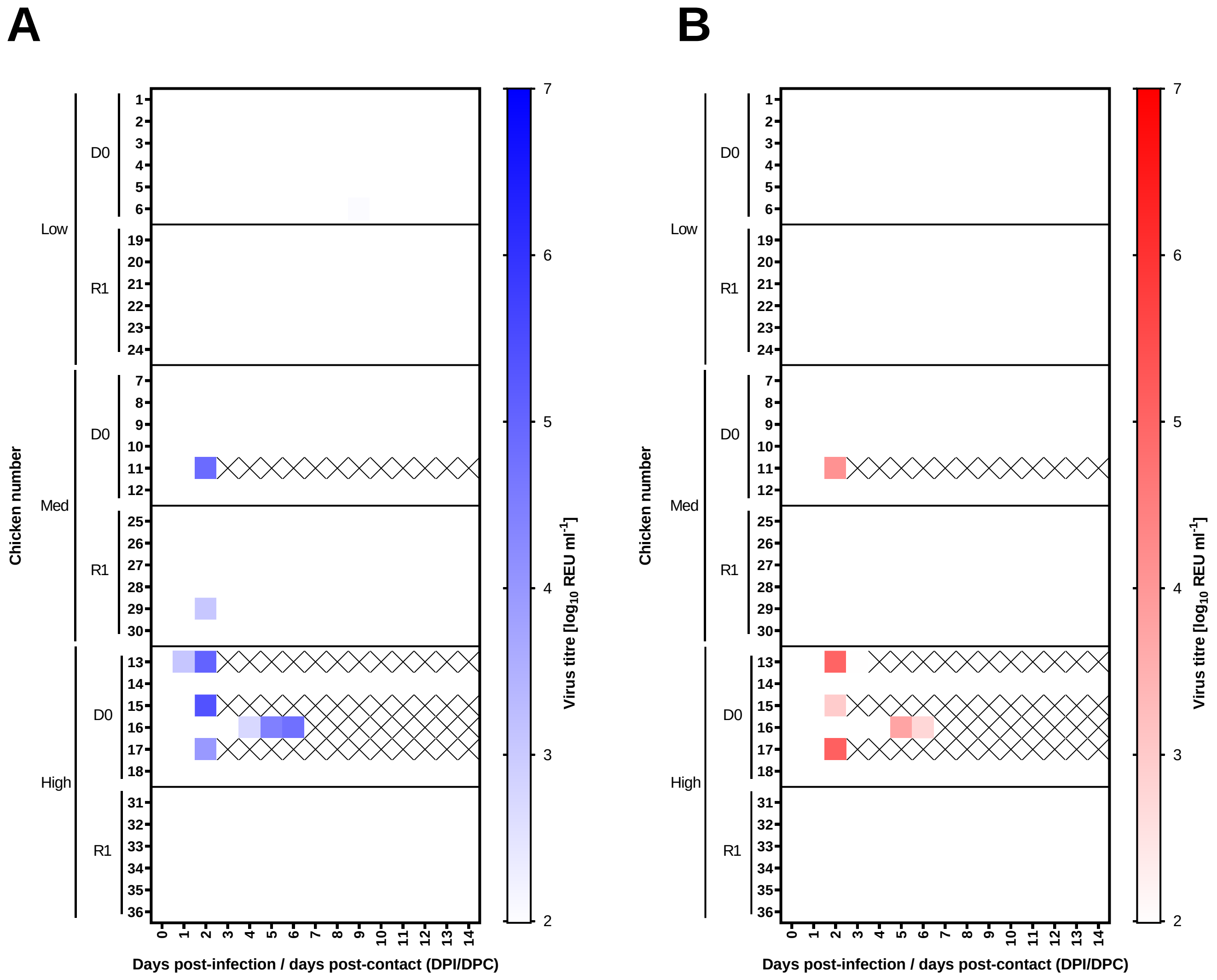


**Fig. S4. Virus shedding from directly infected and contact chickens.**

Virus RNA titres from directly infected (D0) or contact (R1) chickens. D0 chickens were infected with either a low ($10^3$ $EID_{50}$), medium ($10^4$ $EID_{50}$) or high ($10^5$ $EID_{50}$) dose of H5N1-21. Contacts (R1) were introduced 6 hours after infection into the pen of each dose group. vRNA titres derived from swabs taken from the Op (A; blue) or C cavities (B; red). Virus titres were determined using the M-gene RRT-PCR and quantified as REU/ml (see Methods). Viral titres were displayed graphically as a heat map, with white indicating no Ct or REU values below the limit of positivity. Shaded colours representing higher vRNA titres. a 'cross' indicates no sample collected at a specific time point due to the bird being euthanised following severe clinical signs.
